# Centrifugal Gel Crushing for Gel-based Proteome Analysis

**DOI:** 10.1101/2023.06.15.545122

**Authors:** Hiroshi Nishida, Eisuke Kanao, Yasushi Ishihama

**Affiliations:** Division of Medicinal Frontier Sciences, Graduate School of Pharmaceutical Sciences, Kyoto University, Kyoto 606–8501, Japan; Laboratory of Clinical and Analytical Chemistry, National Institute of Biomedical Innovation, Health and Nutrition, Ibaraki, Osaka 567-0085, Japan

**Keywords:** In-gel digestion, SDS-PAGE, Gel crushing, GeLC/MS/MS

## Abstract

We have developed a centrifugal gel crushing method using a pipette tip. Polyacrylamide gel slices are extruded from the narrowing cavity of a pipette tip by centrifugation in a few minutes to crush them into pieces of appropriate size. The size of the crushed gel could be controlled by several parameters, including centrifugal force and pipette tip cavity. In shotgun proteomics, gel-based LC/MS/MS, so-called GeLC/MS/MS, involves the essential but tedious processes of pre-fractionation by SDS-PAGE, followed by dicing the entire gel lane into several parts, fine dicing, and in-gel digestion after the diced gel is manually transferred to a microtube. In this study, we developed an alternative way to crush the pre-fractionated gel slice into small and irregular-shaped gels by centrifugal extrusion of the sliced gel from the narrow cavity of a pipette tip. As a result, we observed an improved recovery and reproducibility of digested proteins compared to the conventional method of manual dicing. We believe that this simple and rapid method of crushing polyacrylamide gels, which allows for parallel operations and automation, is useful for GeLC/MS/MS analysis and applicable to other approaches including top-down proteomics.

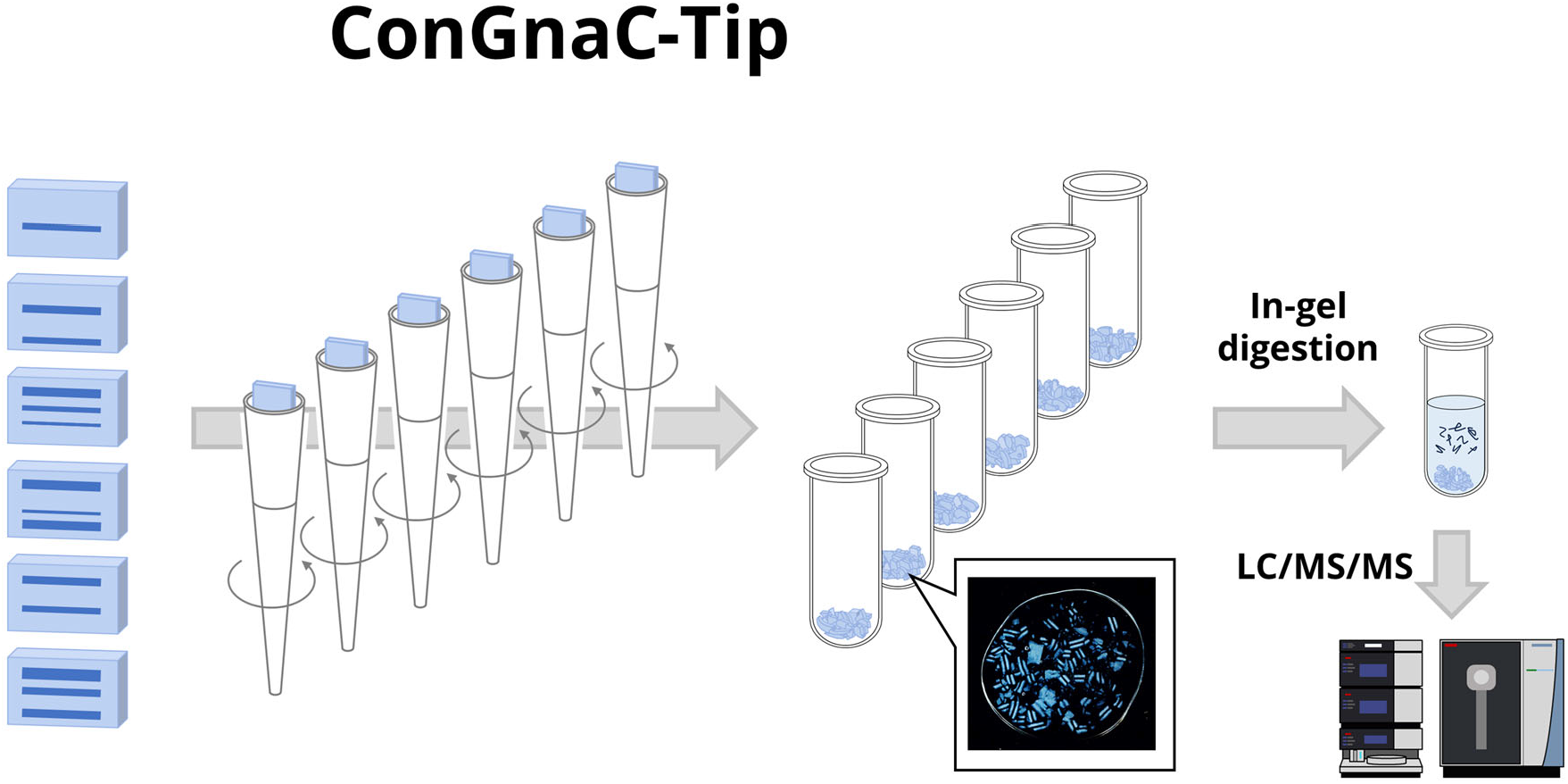

## INTRODUCTION

Polyacrylamide gel electrophoresis (PAGE) including sodium dodecyl sulfate (SDS)-PAGE and isoelectric focusing electrophoresis is a widely used technique for separating proteins at high resolution^1^. In top-down proteomics, SDS-PAGE works to separate and collect proteins of interest according to molecular weight prior to MS analysis ^2,3^, but even in bottom-up proteomics, SDS-PAGE has been used as a pre-fractionator at the protein level, called “GeLCMS’’ approaches ^4,5,6^. Theoretically, protein fractionation should be more effective than peptide fractionation in increasing the number of identical proteins^7^, but GeLCMS approaches require more complex and tedious steps than in-solution digestion-based gel-free LC/MS/MS approaches, and these are difficult to automate and parallelize, so currently GeLCMS approaches are not as widely used as gel-free approaches. One of the key steps in the sample preparation is to cut the gel into cubes of about 1 to 2 mm on each side using a scalpel or razor. This dicing step is performed manually and can be quite time consuming if the number of samples is large, and there is also the possibility of contamination with keratin or other contaminants during the step. Moreover, for such a time-consuming and labor-intensive process, the protein recovery in in-gel digestion is low (approximately 30-50%)^8^.

In this study, we developed a simple and easy method to crush the pre-fractionated gel slice into small and irregular-shaped pieces by centrifugal extrusion of the sliced gel from the narrowing cavity of a plastic pipette tip. We investigated the effect of several parameters, such as centrifugal force and pipette tip cavity, on the size of the crushed gel. In addition, this method was applied to GeLC/MS/MS analysis of protein mixtures extracted from human cultured cells to evaluate its performance.

## EXPERIMENTAL SECTION

### Materials and Reagents

TPCK-treated sequencing-grade modified trypsin was obtained from Promega (Madison, WI). UltraPure™ Tris Buffer was purchased from Thermo Fisher Scientific (Waltham, MA). SDB-XC Empore disks for desalting and their adapters for microtubes were from GL Sciences (Tokyo, Japan). Water was purified by a Millipore Milli-Q system (Bedford, MA). Protease inhibitor cocktail was purchased from Sigma-Aldrich (St. Louis, MO). All other chemicals including s 30 w/v% acrylamide/bis mixed solution (29:1), ammonium persulfate (APS) and *N*-tetrametylethylenediamine (TEMED) were purchased from FUJIFILM Wako (Osaka, Japan) unless otherwise specified. Protein LoBind® microtubes (2 mL) and epT.I.P.S.® pipette tips (1000 µL) were purchased from Eppendorf (Hamburg, Germany).

### Cell Culture and Protein Extraction

HEK293T cells obtained from the RIKEN BRC cell bank (Ibaraki, Japan) were cultured in Dulbecco’s modified Eagle’s medium with 10 % fetal bovine serum and 100 μg/mL Kanamycin at 37 °C, 5 % CO_2_. HEK293T cells were suspended in an SDS lysis buffer consisting of protease inhibitors, 2% SDS, 50 mM Tris-HCl buffer (pH 7.8). The lysate was sonicated, mixed with an equal volume of bromophenol blue solution consisting of 4% SDS, 10% sucrose, 0.004% bromophenol blue and 0.25 M Tris-HCl (pH 6.8) and boiled at 95 °C for 5 min.

### SDS-PAGE and in-gel digestion

We made the polyacrylamide gels by ourselves as follows. First, a 1 mm thickness 10%T, 3.3%C polyacrylamide gel was prepared as a resolving gel: an acrylamide solution consisting of 10 w/v% acrylamide/bis (29:1), 375 mM Tris (pH 8.8) and 0.1 w/v% SDS was mixed with APS and TEMED to a final concentration of 0.83 w/v% and 0.08 v/v%, respectively, poured into assembled glass sandwich plates and allowed to polymerize for approximately 1 hour. Next, a 3 %T, 3.3 %C polyacrylamide gel was then prepared as a stacking gel: an acrylamide solution consisting of 10 w/v% acrylamide/bis, 125 mM Tris (pH 6.8), 0.1 w/v% SDS was mixed with APS and TEMED to a final concentration of 0.83 w/v% and 0.08 v/v%, respectively, poured on top of the resolving gel. A comb was inserted into the unpolymerized stacking gel and allowed to polymerize for at least 1 hour. Electrophoresis was performed by the Mini-PROTEAN® Tetra system purchased from Bio-Rad (Hercules, CA). After electrophoresis, negative staining and destaining were performed according to the instructions of the negative gel stain MS kit from Fujifilm Wako. The gel was diced manually or crushed by centrifugation with pipette tips. For manual dicing, the gel was cut with a scalpel into pieces of 1 ∼ 2 mm per side and transferred into microtubes. For the centrifugal gel crushing, pipette tips were prepared with the tips cut at the appropriate position with a cutter, and gel slices were loaded into the cut pipette tips and extruded into microtubes by centrifugation using an Eppendorf Microcentrifuge model 5415R.

In-gel digestion was performed based on the previously reported protocol^9^. After gel dicing or crushing, reduction was performed by adding 500 μL of 50 mM ammonium bicarbonate buffer containing 10 mM dithiothreitol and sonication for 30 minutes. The reduction solution was then removed and 500 μL of 50 mM ammonium bicarbonate buffer containing 50 mM iodoacetamide was added, and alkylation was performed by sonication in the dark and at room temperature for 30 minutes. Then the gel pieces were washed with 400 μL of methanol/water/acetic acid (MWA, 50:45:5) and sonicated for 30 min. After twice replacements of MWA, the gel pieces were incubated in 500 μL of 50 mM ammonium bicarbonate buffer for 5 min, then in 150 μL of acetonitrile for 15 min, and dried thoroughly in a Speedvac (Thermo Fisher Scientific). The dried small gel pieces were re-swollen in 50 mM ammonium bicarbonate containing 0.01 μg/µL of trypsin, and then 50 μL of 50 mM ammonium bicarbonate was added. The samples were incubated overnight at 37°C. Trypsin-digested peptides were extracted twice by 15 min sonication in acetonitrile/water/trifluoroacetic acid (50:50:0.1) and acetonitrile/water/trifluoroacetic acid (80:20:0.1). The combined extracts were dried in the Speedvac and desalted using StageTips^10,11^.

### NanoLC/MS/MS Analysis

LC/MS/MS analyses were performed on an Ultimate 3000 liquid chromatograph combined with an Orbitrap Exploris 480 mass spectrometer (Thermo Fisher Scientific). The LC mobile phases consisted of solvent A (0.5% acetic acid) and solvent B (0.5% acetic acid and 80% acetonitrile). Peptides were loaded onto self-pulled needle columns (250 mm, 100 μm ID) packed with Reprosil-Pur 120 C18-AQ 1.9 μm (Dr. Maisch, Ammerbuch, Germany)^12^, and separated by a linear gradient set as follows: 5-40 % B in 30 min, 40-99 % B in 0.1 min, followed by 99 % B for 4.9 min. The flow rate was 400 nL/min. The electrospray voltage was set to 2.4 kV in the positive mode. The mass range of the survey scan was from 375 to 1500 m/z with resolution of 60000, 300 % normalized AGC target, and auto maximum injection time. The first mass of MS/MS scan was set to 120 *m/z* with a resolution of 15000, standard automatic gain control, and auto maximum injection time. The fragmentation was performed by higher energy collisional dissociation with a normalized collision energy of 30 %. The dynamic exclusion time was set to 20 s.

### Data Analysis

The raw MS data files acquired by the Orbitrap Exploris 480 mass spectrometer were searched by MSFragger (ver. 3.7) against the UniProtKB database of human proteins containing common contaminants (20437 entries, 2023_02) downloaded via FragPipe (ver 19.1)^13^. The search parameters were loaded with closed search default config, which set two missed cleavage sites, Met oxidation and protein N-terminal acetylation as variable modifications, and Cys carbamidomethylation as a fixed modification. The FDR filter was set to 0.01 at the PSM and protein levels.

### Data Availability

The raw MS data and analysis files have been deposited with the ProteomeXchange Consortium (http://proteomecentral.proteomexchange.org) via the jPOST partner repository (https://jpostdb.org)^14^ with the data set identifier PXD042573.

## RESULTS AND DISCUSSION

### Investigation of parameters for centrifugal gel crushing

We have developed a method for crushing gels using centrifugation and disposable pipette tips. The workflow is depicted in Fig. 1. Briefly, proteins extracted from the sample are separated by SDS-PAGE and the entire lane is sliced into several fractions. The gel slice is loaded into a pipette tip and extruded into the narrowing cavity by centrifugal force. In this process, the gel is torn into small pieces and collected in a tube without loss. This operation can be done in parallel, unlike the manual dicing, and the more gel slices to be crushed, the more efficient it is. We named this method Controlled extrusion of Gel from narrowing Cavity of pipette Tip (ConGnaC-Tip).

**Figure 1.**
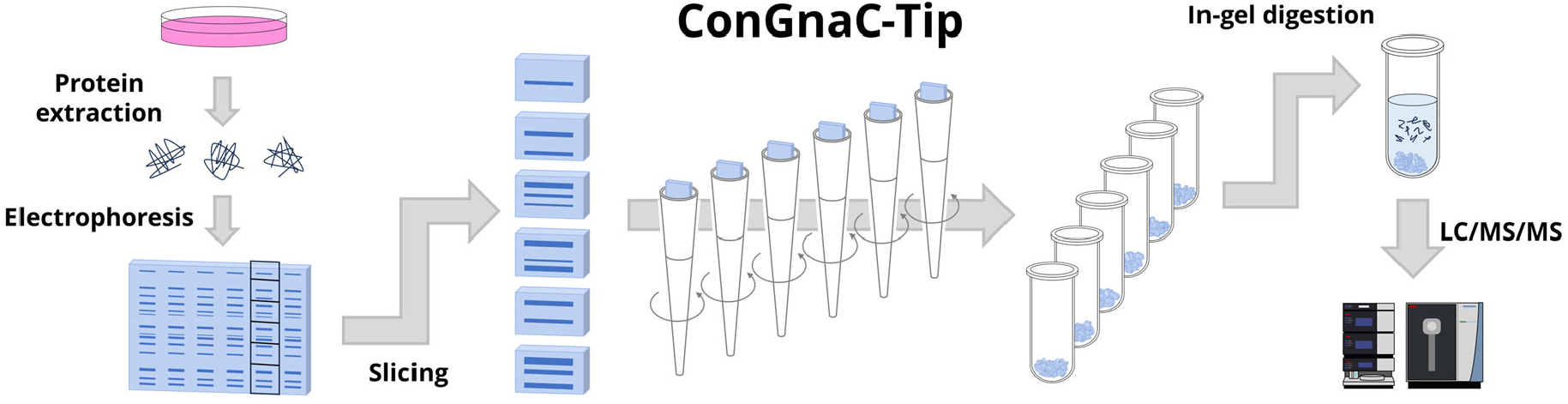
Schematic workflow of gel-based proteome analysis with ConGnaC-Tip. Extracted proteins from the sample were separated by SDS-PAGE followed by slicing and gel crushing using ConGnaC-Tip. After in-gel digestion, the peptides are analyzed by LC/MS/MS.

We characterized the crushed gel size with ConGnaC-Tip when several parameters changed. We used self-made polyacrylamide gels (10%T, 3.3%C) that were not loaded with anything but negatively stained to make it easier to visualize the size. ConGnaC-Tip was performed with plastic pipette tips for 1000 μL, 2 mL microtubes and the microcentrifuge. Adapters for pipette tips were placed on microtubes containing pure water, and polyacrylamide gel was loaded in pipette tips and centrifuged (Figure 1A). First, the effect of the cavity size of pipette tips on the size of the crushed gel was examined using pipette tips cut with a cutter 3, 5, and 7 mm above the tip and intact pipette tips (0 mm). The diameters of the pipette tip cavity cut at 0, 3, 5 and 7 mm were 1.01±0.01, 1.431±0.06, 1.76±0.08 and 2.13±0.02 mm, respectively. The cut gel (5 mm × 10 mm × 1 mm) was loaded in each tip and centrifuged at 5000 x g for 3 min. As a result, the smaller the cavity size of the pipette tip became, the smaller the size of the crushed gel became (Figure 1C-F). Next, the effect of centrifugal force was examined. The gel slices were loaded in pipette tips cut at 5 mm above the tip and centrifuged at 3000, 4000, 5000, 6000 and 7000 x g for 3 min. The result showed that the stronger the centrifugal force became, the smaller the size of the crushed gel became (Figure 1G-K). Finally, we checked the effect of water in microtubes that receive crushed gel. The gel slices were loaded in pipette tips cut at 5 mm above the tip and extruded at 5000 x g for 3 min into microtubes with or without pure water. When the empty microtube received gel, more fine fragments were generated indicating that water in the microtube acted as a cushion, preventing the crushed gel from crushing into smaller pieces (Figure 1E, L). These results indicate that the size of the crushed gel is affected not only by the cavity size of the pipette tip and centrifugal force, but also by the cushioning of the microtubes that receive the gel.

Next, SDS-PAGE analysis was performed on 2.5 μg of protein extracted from HEK293T cells using a polyacrylamide gel (10%T, 3.3%C). After electrophoresis for about 10 minutes (migration distance was about 10 mm), the gel in each lane was manually diced to about 1 mm or crushed at 3000 to 7000 x g with ConGnaC-Tips cut 5 mm above the tip in triplicate. Proteins were then digested, followed by LC/MS/MS analysis. The identification numbers of peptides and proteins with the centrifugal gel crushing by ConGnaC-Tips at 3000-7000 x g were compared to those with manual dicing (Figure 2A, B). The results showed that ConGnaC-Tip and manual dicing were almost equivalent except for the 7000 x g case. Then MS signal intensities were compared for the 4480 peptides commonly identified under these six conditions. As a result, ConGnaC-Tip at 5000 x g was slightly higher than that of manual dicing (Figure 2C). In particular, the signal intensity ratios in the case of ConGnaC-Tip to manual dicing of peptides with more than 11 residues were higher than for peptides with 7-10 residues (Figure 2D). Furthermore, the coefficient of variation of protein quantification values by ConGnaC-Tip at 5000 x g was lower than for manual dicing (Figure 2E). These results indicate that the ConGnaC-Tip method with optimized condition can recover trypsin-digested peptides with higher efficiency and reproducibility than the conventional manual dicing method.

**Figure 2.**
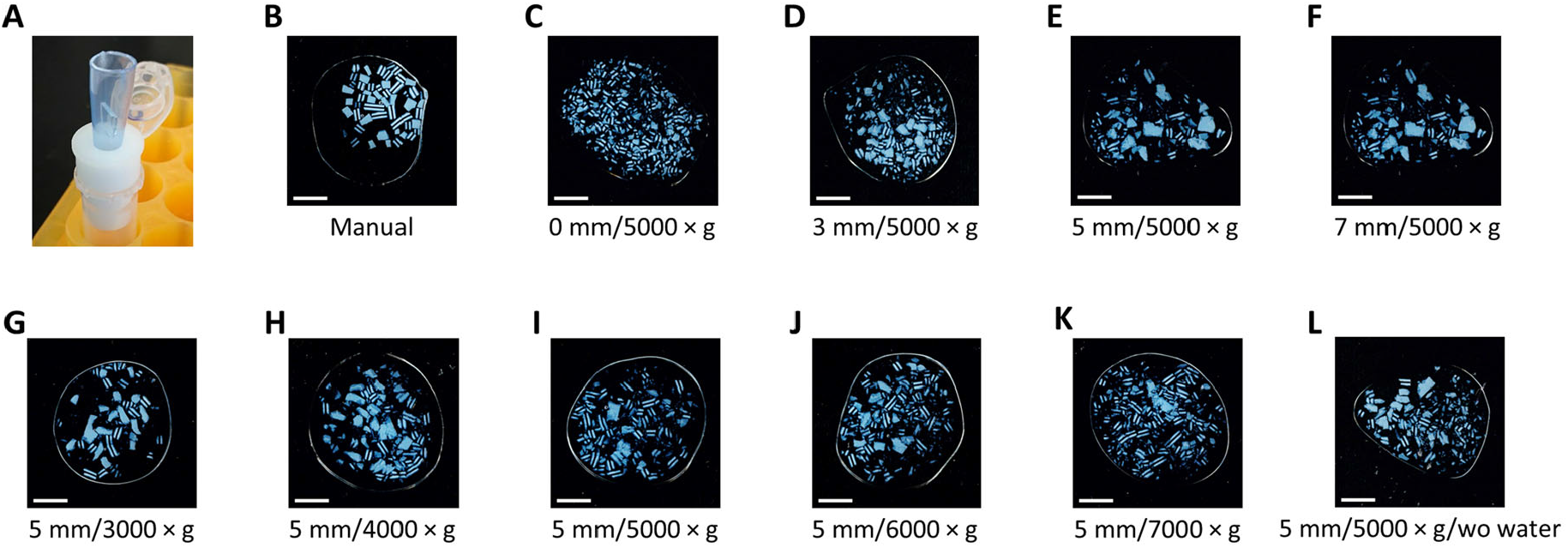
Photographs of ConGnaC-Tip treated gel pieces. (A) Gel was loaded in a 1000 µL-sized pipette tip (with the top cut off to avoid touching a lid of a centrifuge machine), placed on an adapter and centrifuged. Photograph of (B) manually cut or (C-L) centrifugal gel crushing with ConGnaC-Tip under several conditions. The white line in the bottom left corner of each photograph indicates a length of 5 mm. The stripe in the middle of the gel pieces is the unstained area left after negative staining from both sides, and was intentionally left after controlled staining time so that it could be recognized whether it was the surface or the cross-section of the gel.

**Figure 3.**
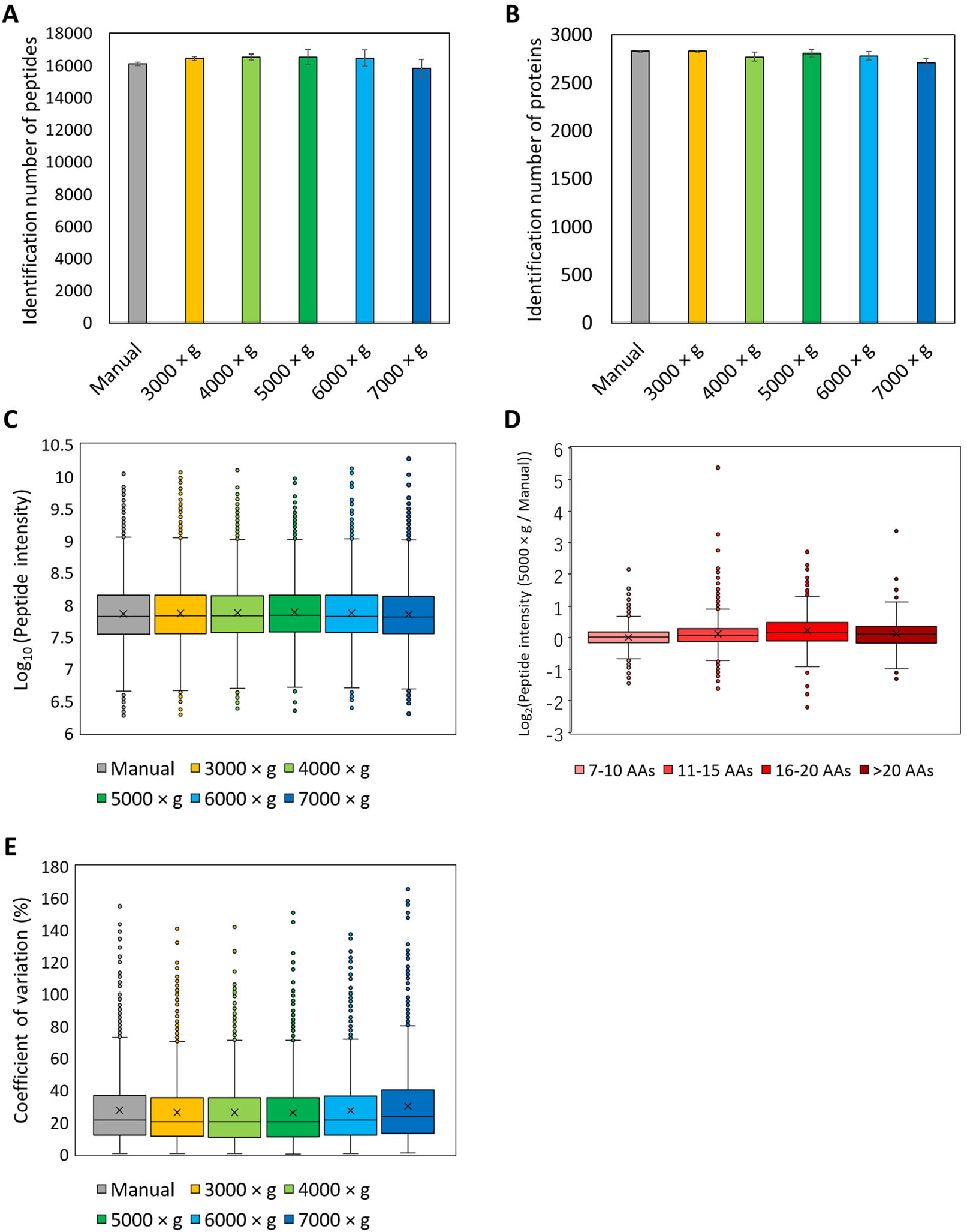
Evaluation of ConGnaC-Tip performance in GeLC/MS/MS analysis of 2.5 µg HEK293T extracted proteins. After SDS-PAGE, the gel pieces were prepared by manual dicing or ConGnaC-Tip with the tip cut 5 mm. Different centrifugation at 3000, 4000, 5000, 6000, or 7000 x g was applied. One tenth of the digested proteins were measured by LC/MS/MS. (A) Identification number of peptides under 6 different conditions, (B) Identification number of proteins, (C) Peak intensity distribution of 4480 peptides commonly identified under these 6 conditions. (D) Peptide length dependence of peak intensity ratios for peptides commonly identified by manual dicing and ConGnaC-Tip at 5000 x g. (E) Distribution of coefficients of variation of quantitation values for 1752 commonly identified proteins under 6 different conditions.

The idea behind this approach is konjac, a traditional Japanese food. It is a hydrogel composed of glucose and mannose made from konjac potato starch. When konjac is cooked to soak up the delicious sauce, it is torn into small pieces by hand rather than cut with a knife to maximize the surface area of the gel, making it uneven in size and cross-section. We thought we could use the same principle to maximize the efficiency of immersion of the trypsin solution into the gel pieces and the extraction of peptides from the gel. The obtained results were as expected, and the developed centrifugal gel crushing method using a pipette tip can be easily automated and parallelized, unlike the tearing konjac, which is torn off one by one by hand.

## CONCLUSION

The ConGnaC-Tip approach was developed to crush polyacrylamide gel slices into small pieces easily and rapidly with only pipette tips and centrifugation. The size of the crushed gel could be adjusted by the cavity size of the pipette tip and the centrifugal force. Peptides could be extracted more efficiently from the gel crushed by pipette tips than those cut manually. The extraction efficiency was better for longer peptides. Note that parameters such as tip cavity size and centrifugal force were optimized under our research environment and may need to be optimized further depending on the type of gel (%T/%C, gel thickness, slice size, etc.) and pipette tip shape. Since the ConGnaC-Tip approach can be automated and parallelized, it is expected to be applied to high-throughput GeLCMS analysis. Furthermore, the ConGnaC-Tip approach is expected to be used not only for shotgun GeLCMS, but also for top-down proteomics, where no digestion is performed after high-resolution protein separation including SDS-PAGE.

## ACKNOWLEDGMENTS

We thank Mr. Takumi Kudo for his technical assistance. We would also like to thank members of the Department of Molecular Systems BioAnalysis for fruitful discussions. H. N. was supported by a fellowship for young scientists from the Japan Society for the Promotion of Science (JSPS). This work was supported by the JST Strategic Basic Research Program CREST (No. JPMJCR1862), AMED-CREST program (No. JP18gm1010010), JSPS Grants-in-Aid for Scientific Research 21J15131 to H.N., 21K14652, 23K13774 to E.K. and 21H02459, 23H04924 to Y.I.

## Notes

### Competing Interest Statement

The authors have declared no competing interest.

### Summary of Updates

One of the authors name was wrong in the previous version. (Wrong) Kanao Eisuke, (Corrected) Eisuke Kanao

